# Automated tumor immunophenotyping predicts clinical benefit from anti-PD-L1 immunotherapy

**DOI:** 10.1101/2023.04.03.535467

**Authors:** Xiao Li, Jeffrey Eastham, Jennifer M. Giltnane, Wei Zou, Andries Zijlstra, Evgeniy Tabatsky, Romain Banchereau, Ching-Wei Chang, Barzin Nabet, Namrata Patil, Luciana Molinero, Steve Chui, Maureen Peterson, Shari Lau, Linda Rangell, Yannick Waumans, Mark Kockx, Darya Orlova, Hartmut Koeppen

## Abstract

**Background:** Cancer immunotherapy has transformed the clinical approach to patients with malignancies as profound benefits can be seen in a subset of patients. To identify this subset, biomarker analyses increasingly focus on phenotypic and functional evaluation of the tumor microenvironment (TME) to determine if density, spatial distribution, and cellular composition of immune cell infiltrates can provide prognostic and/or predictive information. Attempts have been made to develop standardized methods to evaluate immune infiltrates in the routine assessment of certain tumor types; however, broad adoption of this approach in clinical decision-making is still missing.

**Methods:** We developed approaches to categorize solid tumors into “Desert”, “Excluded” and “Inflamed” types according to the spatial distribution of CD8+ immune effector cells to determine the prognostic and/or predictive implications of such labels. To overcome the limitations of this subjective approach we incrementally developed four automated analysis pipelines of increasing granularity and complexity for density and pattern assessment of immune effector cells.

**Results:** We show that categorization based on “manual” observation is predictive for clinical benefit from anti-programmed cell death ligand-1 (PD-L1) therapy in two large cohorts of patients with non-small cell lung cancer (NSCLC) or triple-negative breast cancer (TNBC). For the automated analysis we demonstrate that a combined approach outperforms individual pipelines and successfully relates spatial features to pathologist-based read-outs and patient response to therapy.

**Conclusions:** Our findings suggest tumor immunophenotype (IP) generated by automated analysis pipelines should be evaluated further as potential predictive biomarkers for cancer immunotherapy.

**What is already known on this topic:** Clinical benefit from checkpoint inhibitor-targeted therapies is realized only in a subset of patients. Robust biomarkers to identify patients who may respond to such therapies are needed.

**What this study adds:** We have developed manual and automated approaches to categorize tumors into immunophenotypes based on the spatial distribution of CD8+ T effector cells that predict clinical benefit from anti-PD-L1 immunotherapy for patients with advanced non-small cell lung cancer or triple-negative breast cancer.

**How this study might affect research, practice or policy:** Tumor immunophenotypes should be further validated as predictive biomarker for checkpoint inhibitor-targeted therapies in prospective clinical studies.

## Background

Immuno-oncology (IO) has been practice-changing for the clinical care of patients with a variety of malignancies; long-lasting responses including cures have been observed in patients treated with therapeutics targeting checkpoint inhibitor (CI) molecules such as Cytotoxic T Lymphocyte Antigen 4 and Programmed Death Ligand 1 (PD-L1) [1] [2] [3]. Furthermore, therapeutics targeting other CI molecules or antibodies with immunomodulatory properties such as bi-specific antibodies have shown promise in early stages of clinical evaluation [4]. However, these encouraging results with IO therapeutics are seen only in a subset of patients [5]. This realization has triggered extensive efforts to identify predictive biomarkers in order to maximize efficacy while identifying patients unlikely to respond but who would still be exposed to potentially life-threatening side effects.

Three assays have received regulatory approval for prospective patient identification for CI-targeted therapy: Immunohistochemistry (IHC) for PD-L1 and assessment of tumor mutational burden or microsatellite instability. PD-L1 IHC is by far the most commonly used methodology; however, the requirement of different assays and scoring algorithms across different indications and the imperfect correlation between IHC results and therapy outcome have caused frustration amongst clinicians and pathologists [6] [7] [8] [9] [10].

Success of IO relies on generating or facilitating an anti-tumor immune response. The vast number of known interactions between immune cells (IC) and an established tumor along with the cellular complexity and plasticity of such an immune response makes it difficult to identify a single parameter with sufficient predictive power [11] [12] [13]. Density and phenotype of IC in the TME have previously been used to identify patients with better clinical outcome and/or response to therapy [14]. The Immunoscore [15] [16] [17], originally developed for primary colorectal cancer, evaluates the spatial distribution and density of Cluster of Differentiation 3-positive (CD3+) and CD8+ T lymphocytes and identifies patients with improved clinical outcome independent of prognostic factors such as age, sex, tumor and lymph node status. This approach has been validated in a large patient cohort across multiple institutions using a digital pathology-based image analysis tool [17]. A scoring system for tumor-infiltrating lymphocytes (TILs) has been developed for ductal breast cancer and other carcinomas. It estimates on hematoxylin-eosin (H&E) stained sections the stromal density of all mononuclear cells including plasma cells but excludes granulocytes as well as intra-epithelial immune cells [18] [19] [20] [21]. High density of stromal TILs tends to correlate with better clinical outcome but is also associated with inter-observer variability [22] [23]. Lately, different approaches to assess TILs in solid tumors using digital quantification have been published [24] [25] [26] [27]. Heterogeneity of these approaches with respect to tumor indication, methodology to identify TILs and readouts makes it challenging to compare results and emphasizes the need for standardization as well as validation in large and well-annotated clinical cohorts [28].

Recent studies have described approaches using image analysis with or without machine-learning to delineate the spatial distribution of immune effector cells, both stromal and intra-epithelial, correlating the pattern with gene expression signatures [29] and clinical outcome for patients treated with CI-targeted therapies [30]. Here, we categorized tumors into “Desert”, “Excluded” and “Inflamed” immunophenotypes (IP) based on the density and spatial distribution of CD8+ T effector cells. We reasoned that such categories would provide prognostic and/or predictive information for patients treated with the anti-PD-L1 agent atezolizumab. We show that manual immunophenotyping of tumors is predictive for clinical benefit by CI-targeted therapy in two independent cohorts of two distinct indications. We describe the development and performance of an automated classification pipeline that yields equivalent or superior results compared with the manual methodology. Our results indicate that IP information generated by an automated classifier could represent an important predictive biomarker for IO clinical studies.

## Methods

### Patient cohorts and histology slides

We had access to paraffin embedded sections from three clinical trials. POPLAR (NCT01903993) is a randomized phase 2 study of atezolizumab compared with docetaxel in 287 patients with locally advanced or metastatic NSCLC who failed platinum therapy [31]. OAK (NCT02008227) is a randomized phase 3 trial of 1225 patients with identical design as POPLAR [32]. IMpassion130 (NCT02425891) is a phase 3 randomized clinical study of atezolizumab in combination with nab-paclitaxel (atezolizumab arm) compared with placebo with nab-paclitaxel (placebo arm) for 902 patients with previously untreated TNBC [33]. Tissue sections had to show at least 100 viable invasive tumor cells with associated stroma for immunophenotype evaluation. In the rare case where more than one sample had been submitted for a patient, the sample with the largest tumor area was selected for evaluation. Samples from the same patient with divergent manual tumor IP’s were excluded when considering patient response. This resulted in the evaluation of 745 patient samples for OAK, and 800 for IMpassion130 for manual tumor IP versus automated calls. The study was conducted according to the Declaration of Helsinki and all patients had provided written consent.

### Immunohistochemistry (IHC)

Sections were stained for CD8 and pan-cytokeratin (CK) in a dual chromogenic IHC assay on a Discovery ULTRA platform (Roche Tissue Diagnostics (RTD), Tucson, AZ). Primary antibodies added to the sections were at 3μg/ml (CD8; C8144B) and at 1.6μg/ml (pan-CK; AE1/AE3), respectively (Agilent, Santa Clara, CA). Antigen retrieval was performed with Cell Conditioning Solution 1 (CC1, Cat.# 950-224; RTD) for 64 minutes at 95°C and sections were exposed to primary antibody for 32 minutes at 37°C. Specifically-bound primary antibody was detected using OmniMap anti-mouse HRP (Cat.# 760-4310; RTD) with diaminobenzidine (DAB; CD8) and DISCOVERY Purple (pan-CK; Cat.# 760-229; RTD) followed by counterstain with hematoxylin.

### IP assessment - Manual approach

The tumor area was defined as viable tumor cells with associated stroma. Areas of necrosis or intraluminal aggregates of CD8+ cells were excluded from evaluation. CD8+ cells were detected in essentially all cases; tumors with density of CD8+ cells of “0”, i.e. single dispersed cells with a density too sparse to allow identification of a distinct pattern, were classified as “Desert” (Figure 1A). “Excluded” tumors showed distribution of CD8+ cells limited to the CK- stroma (Figure 1B). “Inflamed” tumors showed co-localization of CD8+ cells with CK+ tumor cells either as a diffuse infiltrate with or without involvement of CK- stroma, or with a predominantly stromal distribution of CD8+ cells with “spill-over” into CK+ tumor cell aggregates (Figure 1C & D). We considered these two patterns as respective ends of a spectrum and did not distinguish between them for our downstream analysis. To address intratumoral heterogeneity of density and pattern of the infiltrate (Figure 1 E & F) we implemented a 20% cut-off; patterns occupying less than 20% of the tumor area were not considered for categorization. Cases showing an inflamed phenotype in >20% of the tumor area were labeled “Inflamed”, independent of the pattern(s) observed in the remaining areas. Implementation of the cutoff was based on a subjective estimate by the inspecting pathologist (Figure 1G).

**Figure 1.**
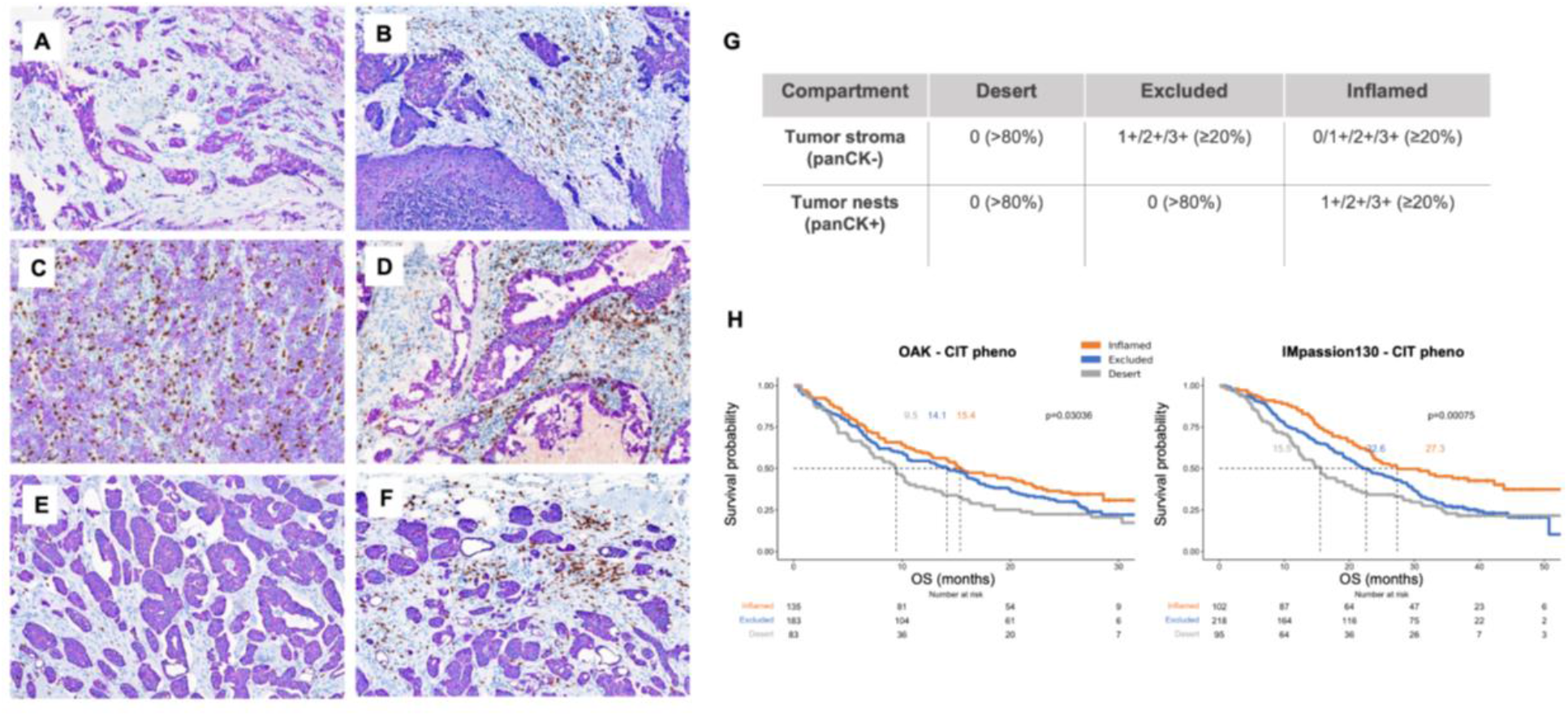
Manually assigned tumor IP and association with treatment outcome. Representative examples of Desert (**A**), Excluded (**B**) and Inflamed (**C** & **D**) immunophenotypes; areas from the same specimen demonstrating areas of Desert (**E**) and Excluded (**F**; intratumoral heterogeneity). Shown are five cases of NSCLC stained for CD8 and panCK. 200x magnification, scale bar indicates 100µm. Cut-off criteria for density and distribution of CD8+ cells for manual IP assignment; % indicate proportion of tumor area involved; 0, 1+, 2+ and 3+ indicate observed densities of CD8+ cells (**G**). Kaplan-Meier curves for OS for OAK and IMpassion130 according to manually assigned IP for patients receiving atezolizumab (**H**).

### IP assessment - Automated approach

#### Whole slide image (WSI) segmentation and tiling

Slides were scanned on a NanoZoomer XR whole slide imager (Hamamatsu, Bridgewater NJ) with a 20x lens, and the images were used to quantify the tumor and IHC positive areas. Annotation of the tumor area was performed manually, excluding areas of assay or scanning artifact and large areas of necrosis. Within these regions, segmentation of CK+ tumor regions and CD8+ DAB pixels were performed by a custom algorithm using Hue, Saturation, and Value (HSV) color thresholding and standard morphological operations in MatLab 2020b (Mathworks, Natick, MA). Images were analyzed using a sliding window of 280×280 pixels and a stride of 200 pixels (0.46 um per pixel) for every analysis except Pipeline #1. CD8 density was quantified at the pixel level in both the CK+ and CK- compartments in each tile, and recorded separately as tile specific measurements, including tile coordinates to enable spatial distribution analysis.

#### Pipeline #1 - Density cutoff

Whole-slide level algorithmic readouts of CD8 density in the CK+ compartment were used to rank order the POPLAR dataset. Using the relative proportions of manual tumor IP calls as a guide, the lowest 20% of these samples were designated as Desert, the next 40% as Excluded, and the highest 40% as Inflamed. The ratio of CD8+ area to CK+ area cutoffs between the classes in POPLAR, 0.00059 CD8+ area / CK+ area between Desert and Excluded, and 0.005 CD8+ area / CK+ area between Excluded and Inflamed, was then used to classify samples in OAK and IMpassion130.

#### Pipeline #2 - Binned CD8 density

For each slide, 10 bins for the density of CD8+ pixels within the CK+ compartment in each tile were empirically defined (with upper bounds of: 0.005, 0.01, 0.02, 0.04, 0.06, 0.08, 0.12, 0.16, 0.2, 1) and the number of tiles in each bin was recorded separately for both compartments. Tiles with less than 25% of their area containing CK+ area were excluded from this count. The same process was performed using the CK- compartment, resulting in 10 additional bins. These two sets of bins are considered entirely separately, and as such it is possible for one tile to contribute to both sets of bin distributions. This resulted in a 20-element vector for each slide, with 10 measurements from each compartment. The vectors from the entire POPLAR dataset were used to train a three class Support Vector Machine (SVM) in Matlab using built-in hyperparameter optimization. This trained SVM was then validated by predicting the phenotype classification for OAK, and tested without further optimization on IMpassion130.

#### Pipeline #3 - randoM fOrest Classifier witH spAtial statistics (MOCHA)

A collection of spatial analytics approaches based on the lattice system was utilized. Firstly, we extracted the center coordinates of the tiles and treated each tile as a spatial unit. Analogous to spatial epidemiology maps, a CK/CD8 prevalence map was created [34], [35]. Specifically, for each tile, the CK+/CD8+ area was calculated, i.e. number of CK+/CD8+ pixels. We further normalized the areas for each tile divided by the sum of CK/CD8 areas across all tiles for the same slide. In addition, we define CK+ tiles as tiles with ratio ≥0.25, where the ratio is calculated as CK+ pixels/(CK+ pixels + CK- pixels) and CK- tiles otherwise. Secondly, based on the prevalence maps, we derived 50 spatial features that either captured the co-localization of CK and CD8 in each slide or spatial distribution of CK or CD8 in each slide, respectively. Some of the features, such as the CD8-CK ratio and Bhattacharyya coefficient in CK+/CK- regions, allow to quantify the CD8 cell density and their colocalization with tumor/stroma cells, and thus could serve as rough proxy for the manual IP (Figure S1A). Collectively, the following set of features was derived:

a. spatial co-localization features of CK and CD8 in CK+ and CK- tiles, respectively: Bhattacharyya coefficient, Morisita-Horn index, Jaccard index, Sorensen index and bivariate Moran’s Index, co-localized Getis-Ord hotspot [36] [37] [38] [39] and CD8-CK area ratio;
b. spatial distribution features of CK and CD8 in CK+ and CK- tiles, respectively: local and global Moran’s and Geary’s C indices [40] and Getis-Ord hotspot [41];

A three-class random forest classifier was then trained based on the entire POPLAR dataset with these 50 features per slide as input. This trained model was then tested by predicting the phenotype classification for OAK and IMpassion130 without further model optimization.

#### Pipeline #4 - randoM fOrest Classifier witH spAtial statistics and BInned CD8 T-cell dEnsity (MOCHA-BITE)

In order to capture the density information as well as the spatial distribution of CK and CD8, respectively, for each slide, we concatenated the features described in Pipeline #2 and Pipeline #3 for each slide. A three-class random forest classifier was then trained based on the entire POPLAR dataset with the combined feature vector per slide as input. This trained model was then tested by predicting the phenotype classification for OAK and IMpassion130 without further model optimization. The five top performing features are shown in Figure S1B.

## Results

### Manual tumor IP classification

We established a methodology to assign tumor IP based on the spatial distribution and density of CD8+ T effector cells on sections stained for CD8 and CK and assigned each case a label of Desert, Excluded or Inflamed. The impact of this categorization on patient outcome was evaluated in the atezolizumab-treatment arms of the OAK and IMpassion130 clinical studies.

Both studies showed longer patient overall survival (OS) when tumors displayed an Inflamed phenotype; patients with Desert tumors had the lowest OS, while patients with Excluded tumors showed intermediate outcome (Figure 1H). Median OS for Inflamed, Excluded and Desert tumors were 15.4, 14.1 and 9.5 months for OAK and 27.3, 22.6 and 15.5 months for IMpassion130, respectively. Patients in the control arm for either trial did not show differences in clinical outcome based on tumor IP (Figure S2).

### Automated tumor IP classification

In order to address subjectivity and lack of scalability of manual tumor IP assignment, we developed an integrated automated classification approach consisting of four distinct pipelines; all pipelines require pre-processed WSI of CD8/CK-stained slides with annotated tumor areas as input (Figure 2A; also see Materials & Methods for details).

**Figure 2.**
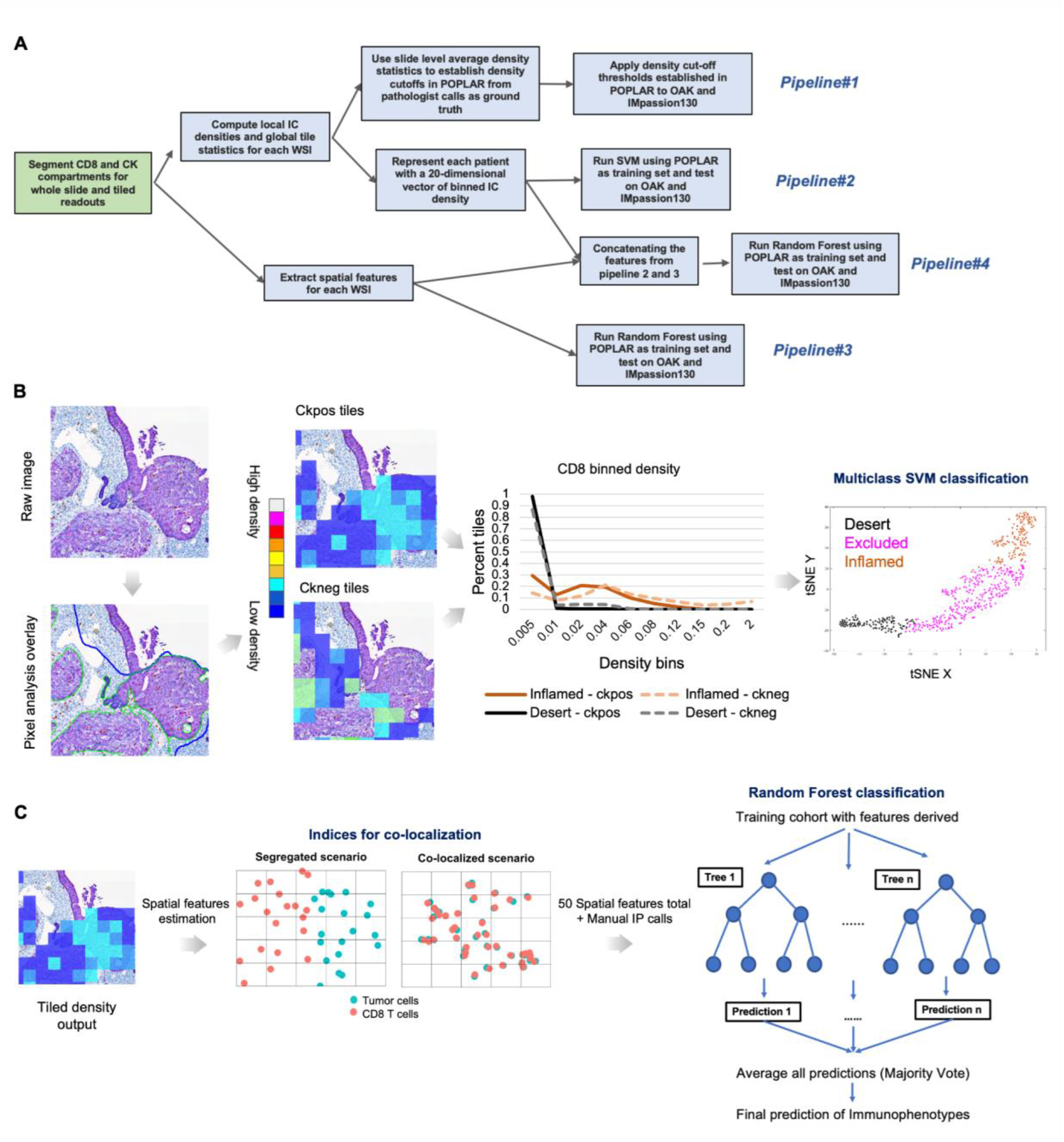
Workflow and output of automated tumor immunophenotype classification pipelines. Input for all four pipelines are whole slide images with manually annotated tumor areas, excluding artifacts (green rectangle; **A**) and manual IP calls from POPLAR. CD8+ and CK+ pixels are automatically identified in the manually annotated tumor area. CD8 density measurements within CK+ regions are the only input for the slide level density cutoff pipeline (#1). CD8 and CK regions are then incorporated in a tile-based analysis, with individual tile densities resulting in a multidimensional readout classified by a SVM into one of three immunophenotype classes, pipeline #2 (**B**). **C**. Data analysis workflow in MOCHA pipeline (#3): based on 50 spatial features (e.g., co-localization of CD8 T cells and tumor cells) extracted for each tiled WSI, a random forest classifier with manual immunophenotype call is trained in a supervised learning framework.

Pipeline#1 is based on the averaged CD8+ cell density score at the slide level as the simplest representation of manually derived IP. In this analysis, the average density of CD8 in the CK+ compartment of the annotated tumor region is determined. We observed a unimodal distribution of CD8 density across all samples with an enrichment of manually classified Desert samples at the low and Inflamed at the high end (Figure 3A). Unimodality of this distribution suggested that fully unsupervised classification of samples into three IP groups would lead to a high misclassification error. IP classification was achieved in a supervised fashion by application of CD8 density cutoffs established using the manual IP calls from POPLAR. The prevalence of manual IP calls in this trial was used to determine these cutoffs, with the 20th percentile of CD8 densities in CK+ regions corresponding to the percentage of Desert samples, resulting in a CD8 density ratio cutoff of 0.059, and the 60th percentile and a CD8 density cutoff of 0.5 for Excluded samples, with the remaining cases assigned to Inflamed samples. These CD8 density cutoffs were then applied to OAK and IMpassion130. A surprisingly high accuracy and linear weighted Cohen’s kappa (0.558 in OAK, and 0.443 in IMpassion130) were observed (Figure 4A, 4C).

**Figure 3.**
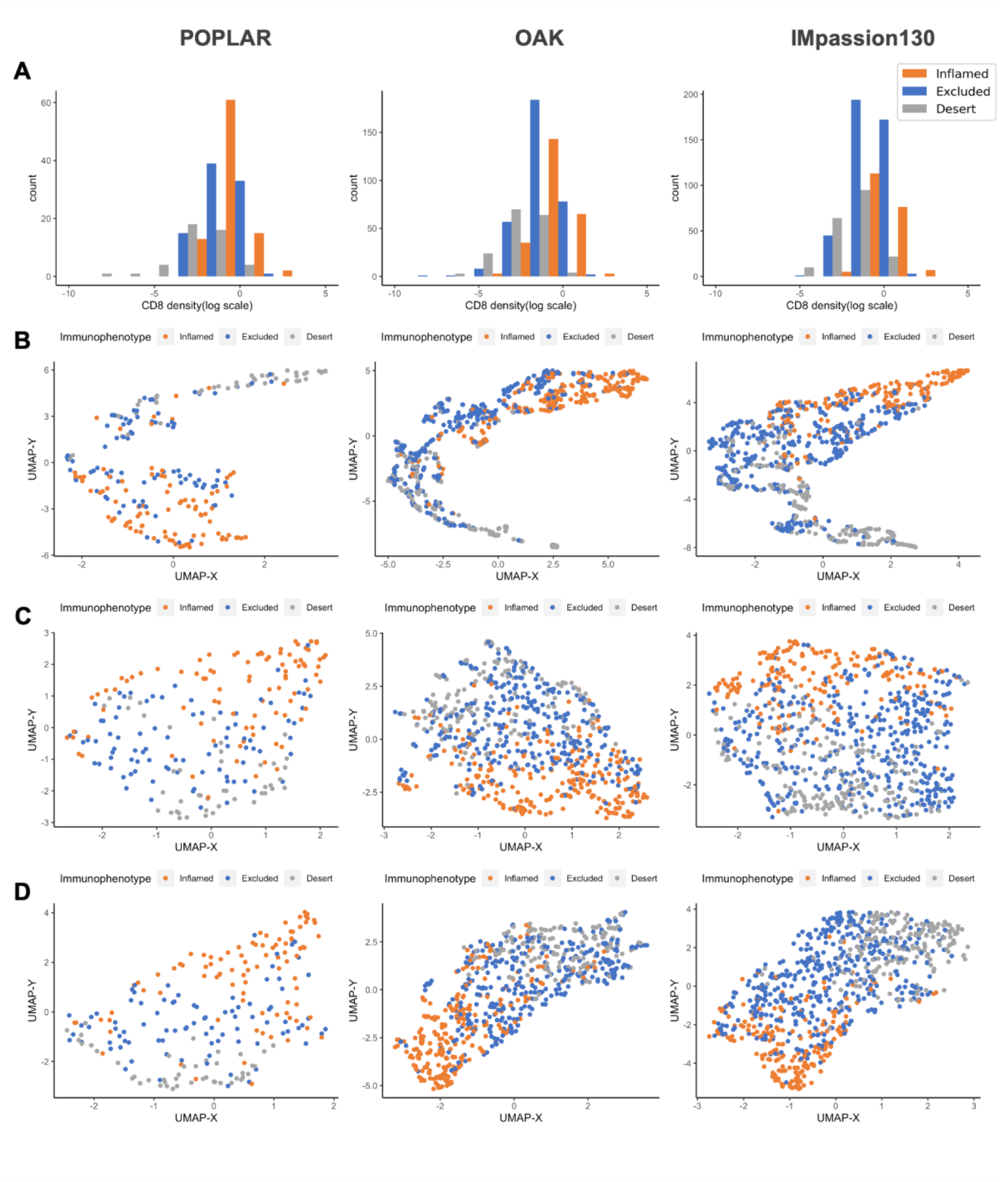
Overlay of the manual tumor IP calls with the features generated by the respective automated analysis pipelines for POPLAR, OAK and IMpassion130. Distribution of patient samples based on the averaged slide level CD8 density readout in CK+ regions (pipeline #1); counts (y-axis) refer to the number of unique patient samples (A). UMAP plots based on: Binned CD8+ cell density in CK+ and CK- tiles (pipeline #2; B); a set of spatial features derived from tile based measurements (pipeline #3; C); a combination of spatial features with binned tile-based CD8 density (pipeline #4; D). Cases are color-coded based on manually assigned tumor IP categories.

**Figure 4.**
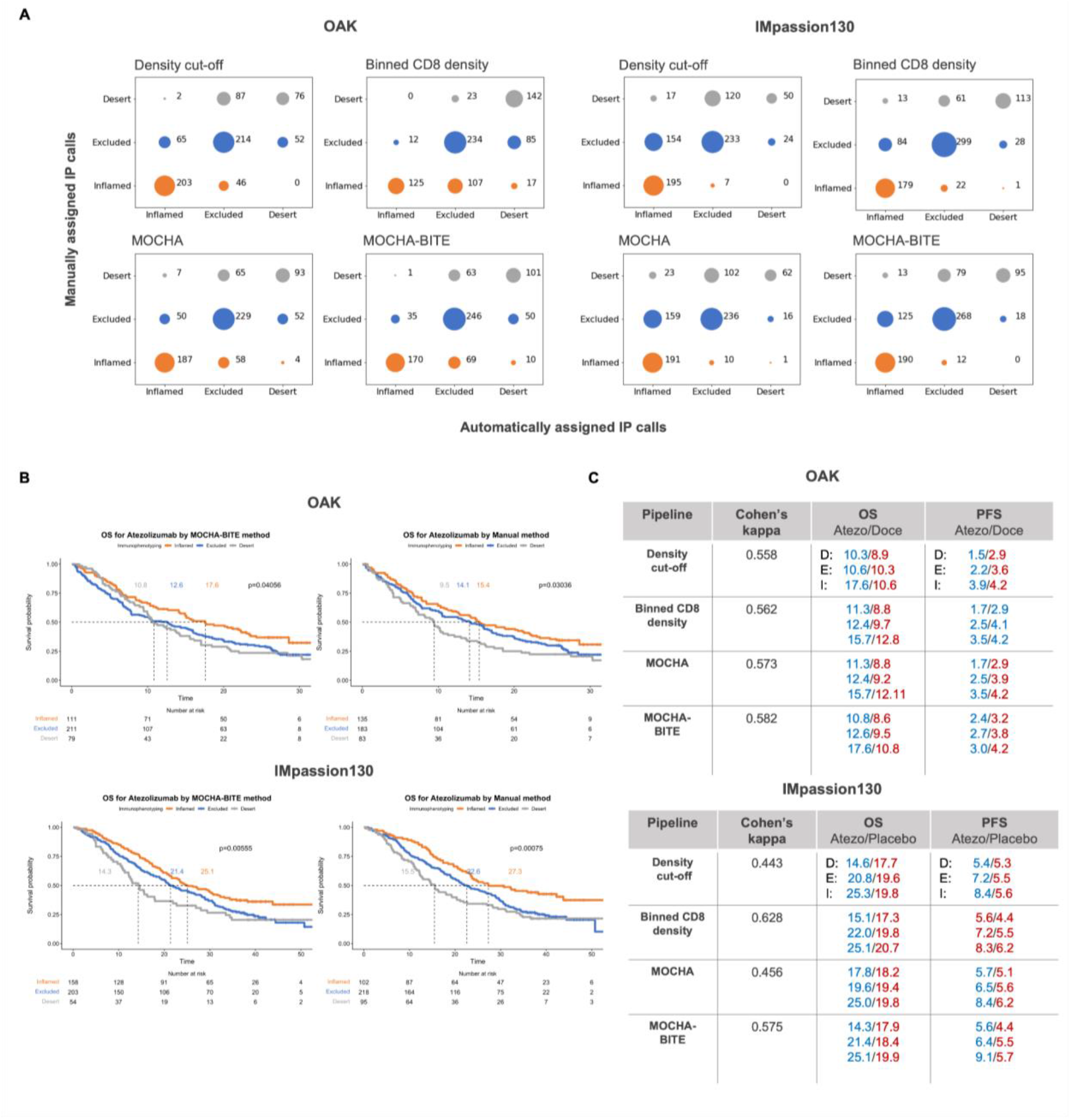
Concordance between manually and algorithmically assigned tumor IP calls and association with treatment outcome. Confusion matrices showing the distribution of manually assigned tumor IP calls (rows) among the automated IP calls (columns) for the four automated pipelines for OAK and IMpassion130 (**A**). Kaplan-Meier curves for OS for atezolizumab-containing treatment arms for OAK and IMpassion130 according to IP categories determined by MOCHA-BITE or manually (see suppl. Fig for PFS curves) **(B)**. Performance of the four automated pipelines based on agreement of IP classification using manual IP calls as ground truth; listed for each pipeline are the Cohen’s kappa coefficient as well as median OS and PFS times for Desert (D), Excluded (E) and Inflamed (I) IP in descending order for the atezolizumab-containing (blue) and control arms (red) (**C**). Blue and red values indicate statistically significant and non-significant differences, respectively according to the log-rank test, separation of OS (or PFS) Kaplan-Meier curves with the given IP categorization.

This same cutoff methodology of using the 20th percentile as the cutoff between Desert and Excluded, and the 60th percentile between Excluded and Inflamed was applied to CD8 density calculated across the entire tumor area without regard to CK+ regions. This resulted in a CD8 density ratio cutoff of 0.003 for Desert samples, and a cutoff of 0.0014 for Excluded samples. This resulted in a decrease in overall accuracy and Cohen’s kappa by approximately 4% for each measure compared to the analysis of the CK+ compartment (data not shown).

Pipeline #2 incorporates the concept of local density used in the manual process. Each image was analyzed using small tiles of 16,000 um^2, providing localized measurements of CD8 density with respect to CK staining (Figure 2B). Within each tile, the CD8+ density was measured separately within CK+ and CK- compartments. The readouts for each tile were divided into 10 bins based on CD8 density within their respective CK+ or CK- compartments, resulting in 20 readouts per sample. These readouts alone, used in the fully unsupervised manner, could not clearly segregate tumors into three clean clusters (Figure 3B); however, there appeared to be a trend for Desert, Excluded and Inflamed enriched regions in that 20 dimensional binned IC density feature space projected into the two dimensional Uniform Manifold Approximation and Projection (UMAP) space. To arrive at a cleaner IP assignment we adopted a supervised classification approach. Each of the twenty CD8+ density bins was used to define a multidimensional feature space, which was separated by a multiclass Support Vector Machine (SVM) trained with the corresponding manual phenotype classifications in POPLAR. The resulting SVM was then used to classify OAK and IMpassion130 (Figure 4A). The weighted Cohen’s kappa for the Binned CD8 density pipeline was 0.562 in OAK, and 0.628 in IMpassion130 compared to the manual immunophenotype classification (Figure 4C).

Pipeline #3 (MOCHA) considers the more granular spatial pattern and density of immune infiltrates (Figure 2A). CK and CD8+ densities calculated in each tile allowed us to obtain a two dimensional density distribution (Figure 2C) of tumor cells and CD8+ T cells. In order to quantitate spatial patterns of tumor cells and CD8 T cells such as cell colocalization, presence of cell “hot spots” etc., we used a collection of lattice-based spatial analytics approaches (see Methods for details). As a result, a set of fifty spatial features was extracted for each tiled WSI from POPLAR, OAK and IMpassion130 datasets. Since these spatial features alone do not clearly differentiate samples into the three IP categories (Figure 3C), we used manual IP calls from the entire POPLAR dataset to train the random forest classifiers and then tested them on OAK and IMpassion130 datasets. This pipeline successfully relates spatial features to manual IP calls (Figure 4A) with the weighted Cohen’s kappa of 0.573 for OAK, and 0.456 for IMpassion130 (Figure 4C).

Pipeline #4 (MOCHA-BITE) combines the 50 spatial features of pipeline #3 with the binned CD8+ cell densities of pipeline #2 and a random forest classifier with manual IP calls is trained in a supervised learning framework. We used IP classification agreement and OS/PFS log-rank test results as criteria to assess pipeline performance.

Based on Cohen’s kappa MOCHA-BITE outperforms the other three pipelines (Figure 4, Figures S1-7) and recapitulates the results observed by manual IP assignment; the only exception is the analysis using pipeline #2 (binned CD8+ cell density) for IMpassion130, which on the other hand performs inferiorly in the PFS analysis (Figure 4C). MOCHA-BITE yielded significant log-rank test results when comparing the three predicted IP groups for OS and PFS using OAK and IMpassion130 as test sets (Figure 4C).

In order to determine if the treatment effect differs between IP patient subgroups (identified by manual or MOCHA-BITE analysis) we performed an interaction test as illustrated in the forest plot analyses for both clinical trials; we focused on OS and “condensed” Desert and Excluded tumors into a “Non-Inflamed” group. We observed that atezolizumab is linked to improved OS clinical benefit in Inflamed vs Non-Inflamed tumors in both, OAK (HR 0.65 [CI 95% 0.48-0.97] vs 0.88 [CI 95% 0.72-1.08], respectively) and IMpassion130 (HR 0.53 [CI 95% 0.38-0.75] vs 0.93 [CI 95% 0.77-1.11], respectively). The difference in HR between Inflamed and Non-Inflamed tumors for either IP classification methodology is statistically significant for IMpassion130 (Figure 5A, 5B). This is also shown in the separation of the accompanying OS KM curves for Inflamed and Non-Inflamed tumors in the atezolizumab vs control arm of each trial.

**Figure 5.**
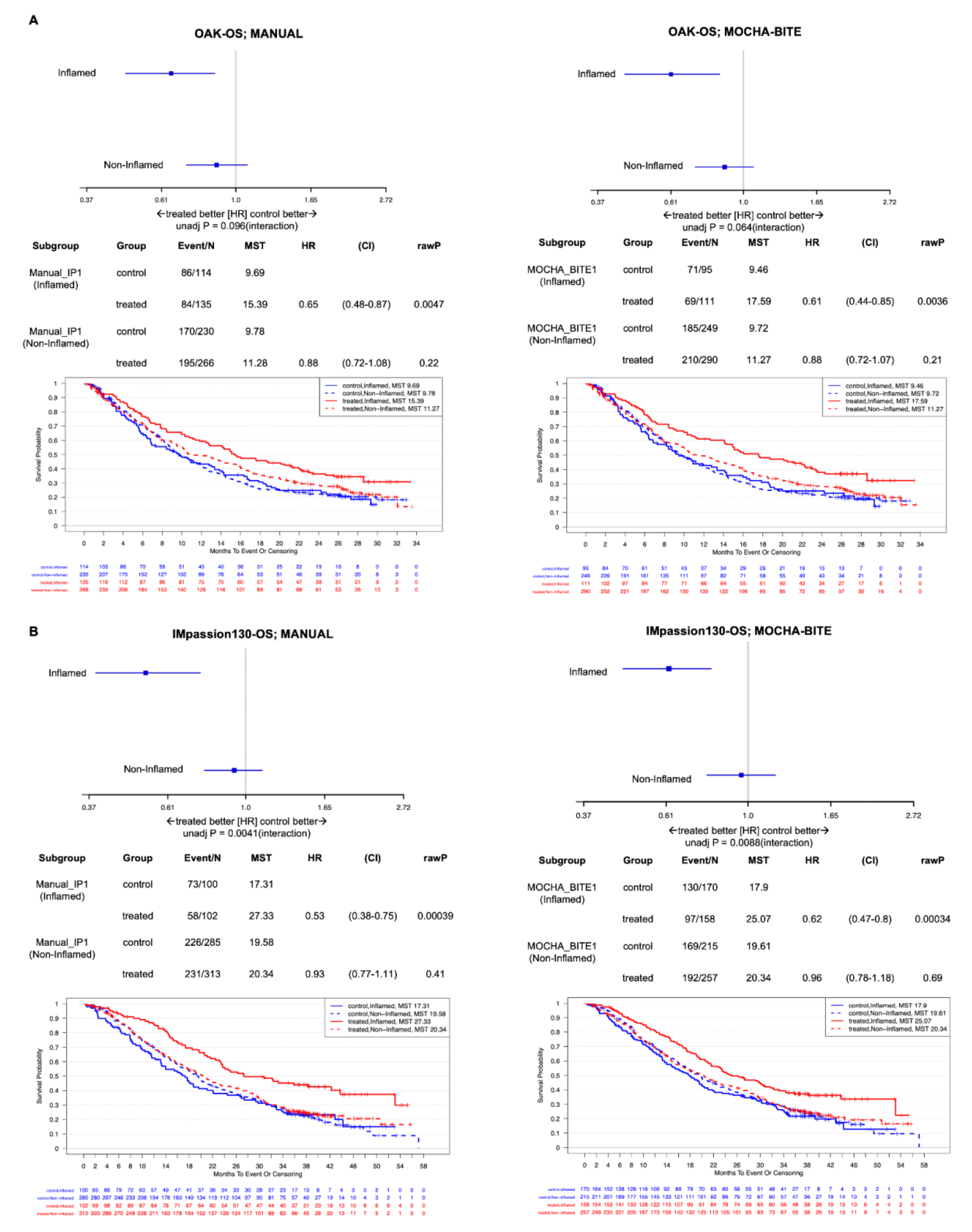
Association of IP with atezolizumab outcome in NSCLC and TNBC patients. Forest plot analysis and Kaplan-Meier curves for patients with “Inflamed” vs “Non-Inflamed” (Excluded+Desert) tumors for OAK (**A**) and IMpassion130 (**B**). The vertical line of no effect in the forest plots indicates a hazard ratio (HR) of 1, shown are the number of patients in control and treatment group for each IP category, median survival times (MST) with confidence intervals (CI) and p values.

## Discussion

Analysis of digital pathology images based on spatial statistics has previously shown value in different clinical use cases for solid and liquid tumors [36] [37] [42] [43] [44] [27]. In this study, we show that tumor IP assignment using different methodologies is predictive for outcome of anti-PD-L1 targeted therapy. We developed our own approach instead of applying previously published algorithms such as Immunoscore [45] or traditional TIL scoring [19] for several reasons. We wanted to pursue the task in an indication-agnostic fashion. Unambiguous identification of epithelial and stromal compartments as well as labeling of the relevant immune cell type allowed us to implement our analyses in two different epithelial tumor types.

Furthermore, diverse specimen types are typically encountered in the clinical trial setting (resections, excisional and needle core biopsies); while our methodology has a minimum requirement of viable, invasive tumor and associated stroma, we did not want to solely rely on resection specimens with representation of tumor center and invasive edge. This would severely limit the number of available specimens and may introduce unwanted bias toward better clinical outcome as resections in NSCLC patients are primarily performed with curative intent. Lastly, we felt strongly about the value of our focus on intra-epithelial CD8+ T-cells in our assessment.

In order to address the limitations of the manual approach of scalability and inter-/intra-observer variability, we developed individual automated pipelines for IP calls and show that using a combined approach (pipeline #4) outperforms individual pipelines in two distinct tumor types.While the training set was limited to cases of NSCLC, implementation of the analysis was successful in TNBC without the need for additional machine-learning efforts. Extension to other tumor types should be straightforward as long as the malignant cells can be unambiguously identified. Furthermore, our approach lends itself to evaluate the distribution of immune cells other than CD8+ T-effector cells as long as they can be identified by immunohistochemical means. This could be relevant for IO therapies targeting molecules other than PD-1 and PD-L1. Also, expansion into a multiplexed methodology with identification of more than one immune cell phenotype should be rather seamless.

An end-to-end deep learning–based Al model that segments cancer epithelium, cancer stroma, as well as TILs to categorize tumors into IP similar to our classification has recently been developed for NSCLC [46]. Similar to our findings, the authors describe improved clinical outcomes for patients with inflamed tumors when treated with anti-PD-1 or anti-PD-L1 agents. The proportion of the three IP categories are similar but far from identical between their and our methodology. There are several reasons for this “discrepancy”: The composition of the respective patient cohorts is most likely different with respect to clinical parameters, cut-offs to define IP categories are different between the two methodologies, and identification of the two tumor compartments and immune cells relies on machine-learning (and thus requires training data) in one and presence/absence of an immunohistochemical stain in the other (our) approach. The latter is an important distinction as it offers the opportunity to expand this approach to other epithelial tumor types without the need to train a new algorithm for recognition of tumor cells in a separate indication. Importantly, two distinct approaches on different patient cohorts show that the spatial distribution of immune cells has predictive value for CI-based therapies suggesting biologically relevant mechanisms that deserve further exploration.

While our results are encouraging in terms of the overall correlation between the immune status of the tumor and patient outcome, there are still the outliers; we observed a certain degree of tumor IP heterogeneity among the patients who responded well to immunotherapy treatment. Incorporating clinical response data in the training of the model may optimize performance and identify features that could serve as additional biomarkers. However, we are also keenly aware of the inherent caveat of the approach of interrogating an environment of high plasticity such as the TME in archival tissues to predict therapy outcome. There is ample clinical and experimental evidence that tumor IP can change during immunotherapy [47] [48] [49] [50]; in vivo imaging approaches could circumvent the need for on-treatment biopsies for immune profiling [51].

We realize that our study – albeit on rather large patient cohorts – is based on retrospective analyses. Further optimization and validation in a prospective setting are necessary prior to routine clinical implementation.

## Ethics approval and consent to participate

All patients provided informed consent to participate in the respective clinical trials.

## Consent for publication

All authors agree to the publication of this manuscript.

## Availability of data and material

Data and materials are available upon reasonable request Code will be made available upon publication

## Contributors

X.L., J.E., D.O. and H.K contributed to the study design and wrote the paper. X.L., J.E., R.B., C.W.C., L.M., B.N., M.K., S.L., L.R., D.O., H.K. made critical contributions to the acquisition and/or analysis of data. All authors were involved in data interpretation and review of the manuscript.

## Acknowledgements

We thank the patients, their families, and all of the investigators and their staff involved in the POPLAR, OAK and IMpassion130 studies. The authors would like to thank the Digital Pathology and Image Acquisition staff and the Histology Archive (both Genentech, Inc.) for scanning of slides, image acquisition and slide retrieval and management, respectively. We would like to thank Robin Lorenz, Lisa McGinnis and Frank Peale for critical review of the manuscript and Meredith Triplett for excellent administrative support.

## List of abbreviations

TME: tumor microenvironment
PD-L1: programmed death ligand-1
NSCLC: non-small cell lung cancer
TNBC: triple-negative breast cancer
IO: immuno-oncology
CI: checkpoint inhibitor
IP: immunophenotype
IHC: immunohistochemistry
H&E: hematoxylin and eosin
IC: immune cells
TILs: tumor-infiltrating lymphocytes
CK: cytokeratin
OS: overall survival
PFS: progression-free survival
WSI: whole slide images
MOCHA: randoM fOrest Classifier witH spAtial statistics
MOCHA-BITE: randoM fOrest Classifier witH spAtial statistics and BInned CD8 T-cell dEnsity

## Supplementary materials

**Supplementary Figure 1.**
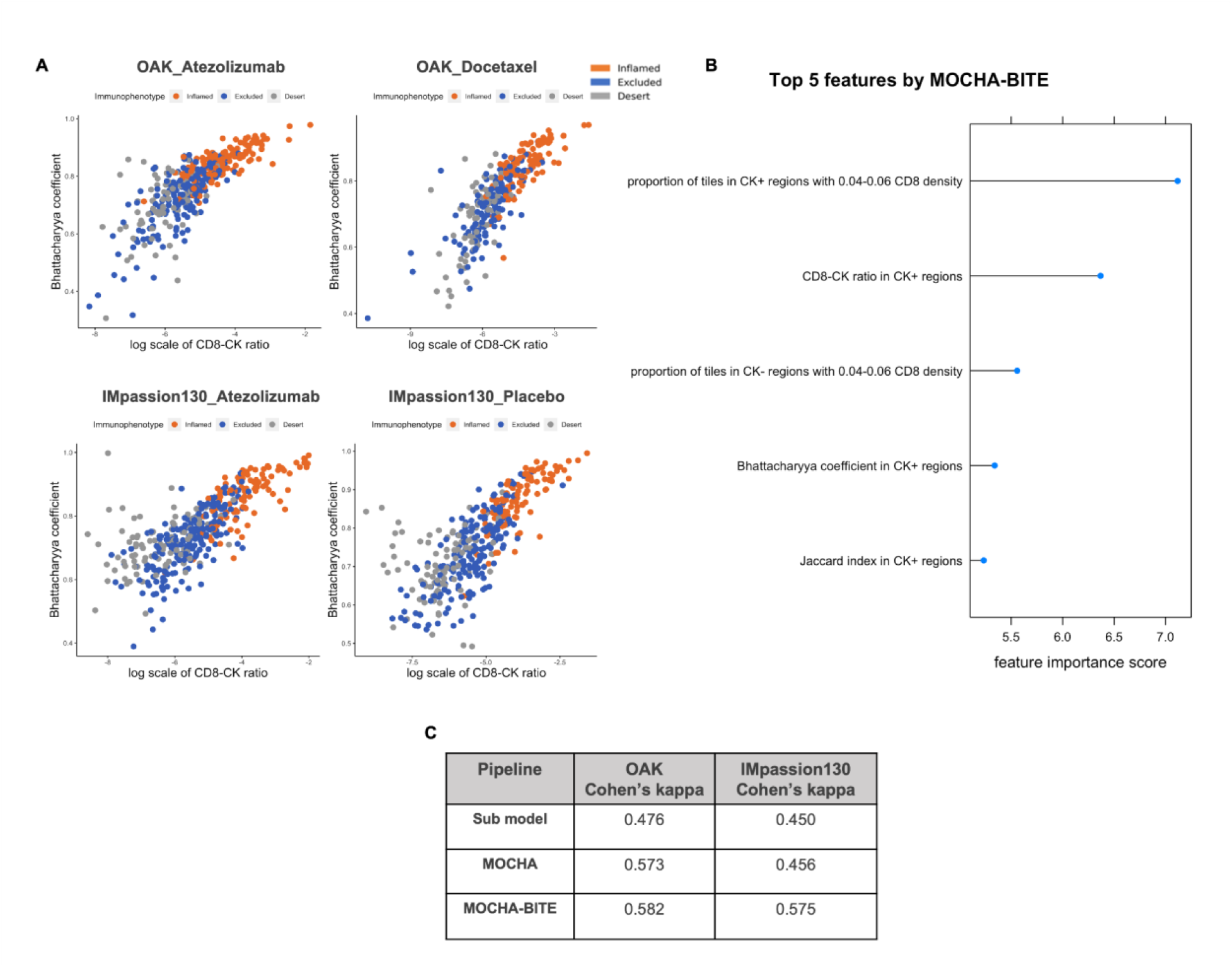
Features extracted in the automated pipelines reasonably approximate the manually assigned tumor IP. MOCHA-BITE pipeline features, such as CD8-CK ratio in CK+ regions and Bhattacharyya coefficient (BC), represent the manual tumor IP well. Each dot corresponds to a unique patient sample. Manual tumor IP category is represented using color-code (**A**). Top five out of fifty total MOCHA-BITE features ranked according to their contribution towards the IP classes separation using POPLAR as a training data set (**B**). Model performance using only CD8-CK ratio in CK+ regions and BC as features (Sub-model) versus MOCHA and MOCHA-BITE (**C**).

**Supplementary Figure 2.**
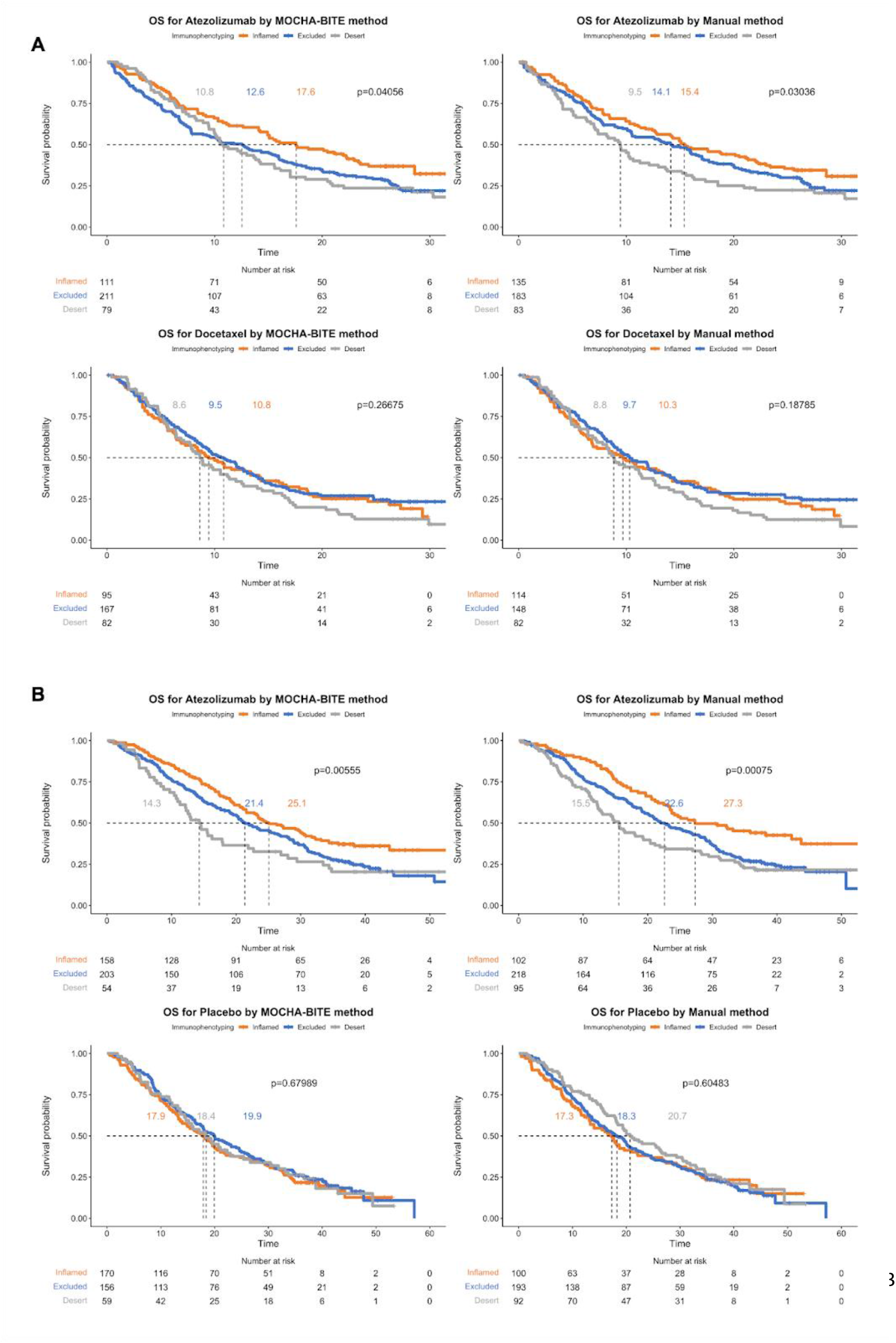
OS log-rank tests for tumor IP subgroups, as classified by MOCHA-BITE and manual methods. Kaplan-Meier curves for OS for the atezolizumab and control treatment arms for OAK **(A)** and IMpassion130 **(B)** with median survival times for the three IP categories as determined by MOCHA-BITE or manually.

**Supplementary Figure 3.**
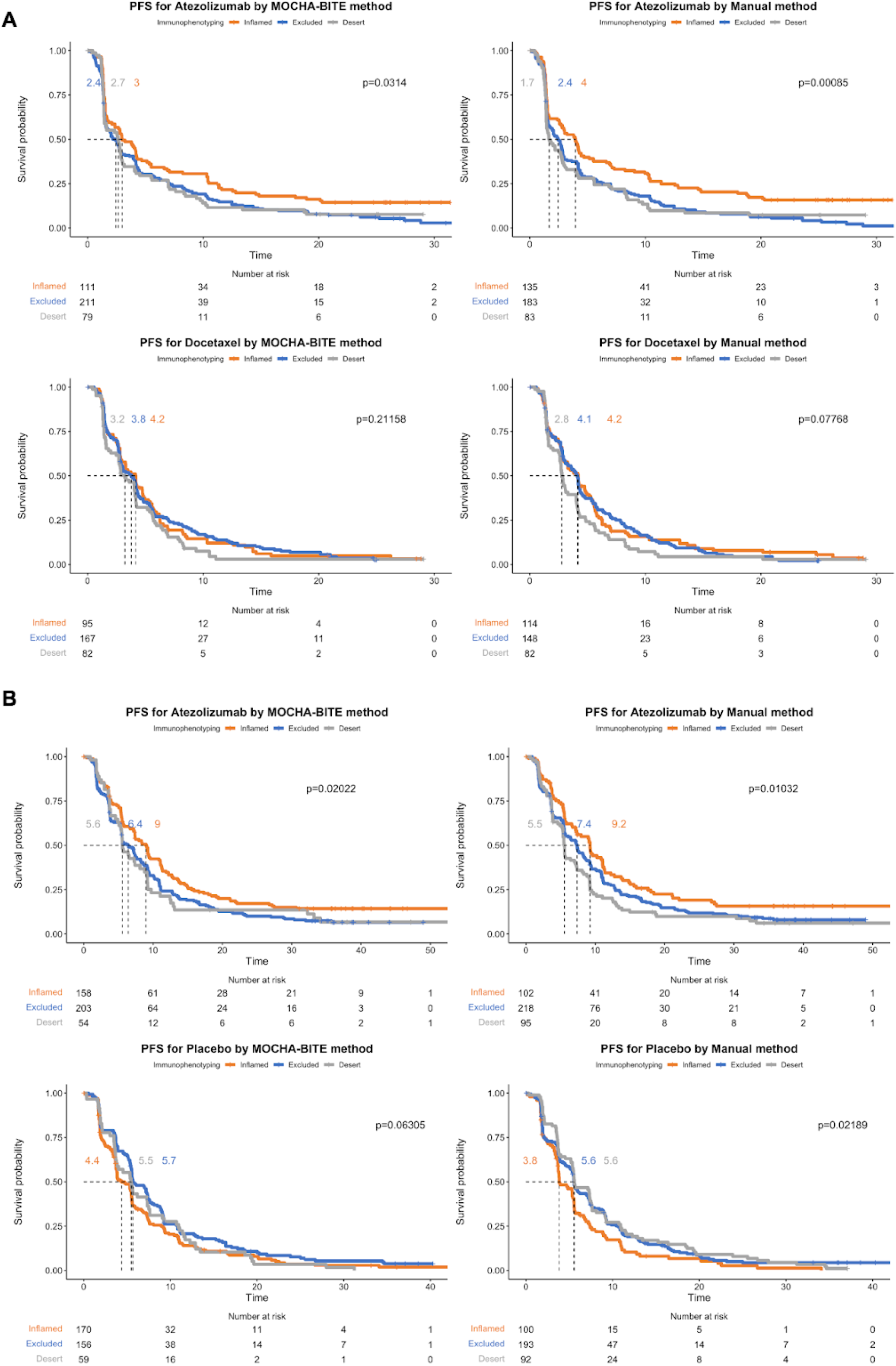
PFS log-rank tests for tumor IP subgroups, as classified by MOCHA-BITE and manual methods. Kaplan-Meier curves for PFS for the atezolizumab and control treatment arms for OAK **(A)** and IMpassion130 **(B)** with median survival times for the three IP categories as determined by MOCHA-BITE or manually.

**Supplementary Figure 4.**
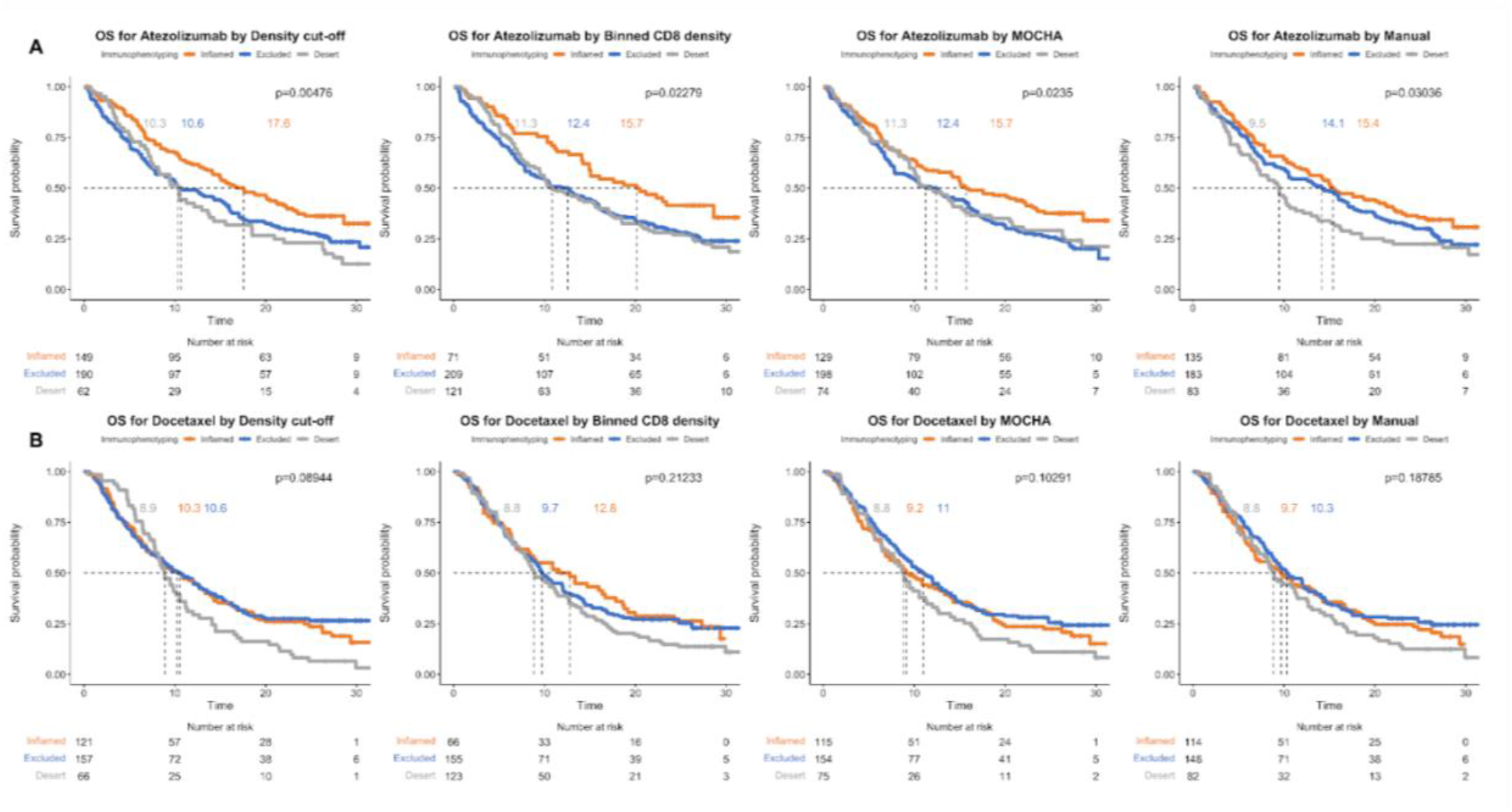
Kaplan–Meier curves for OS in atezolizumab and docetaxel arms in OAK. Log-rank p-values are shown for tumor IP classes identified by the automated pipelines #1-3 and the manual method for atezolizumab arm (**A**) and docetaxel arm (**B**).

**Supplementary Figure 5.**
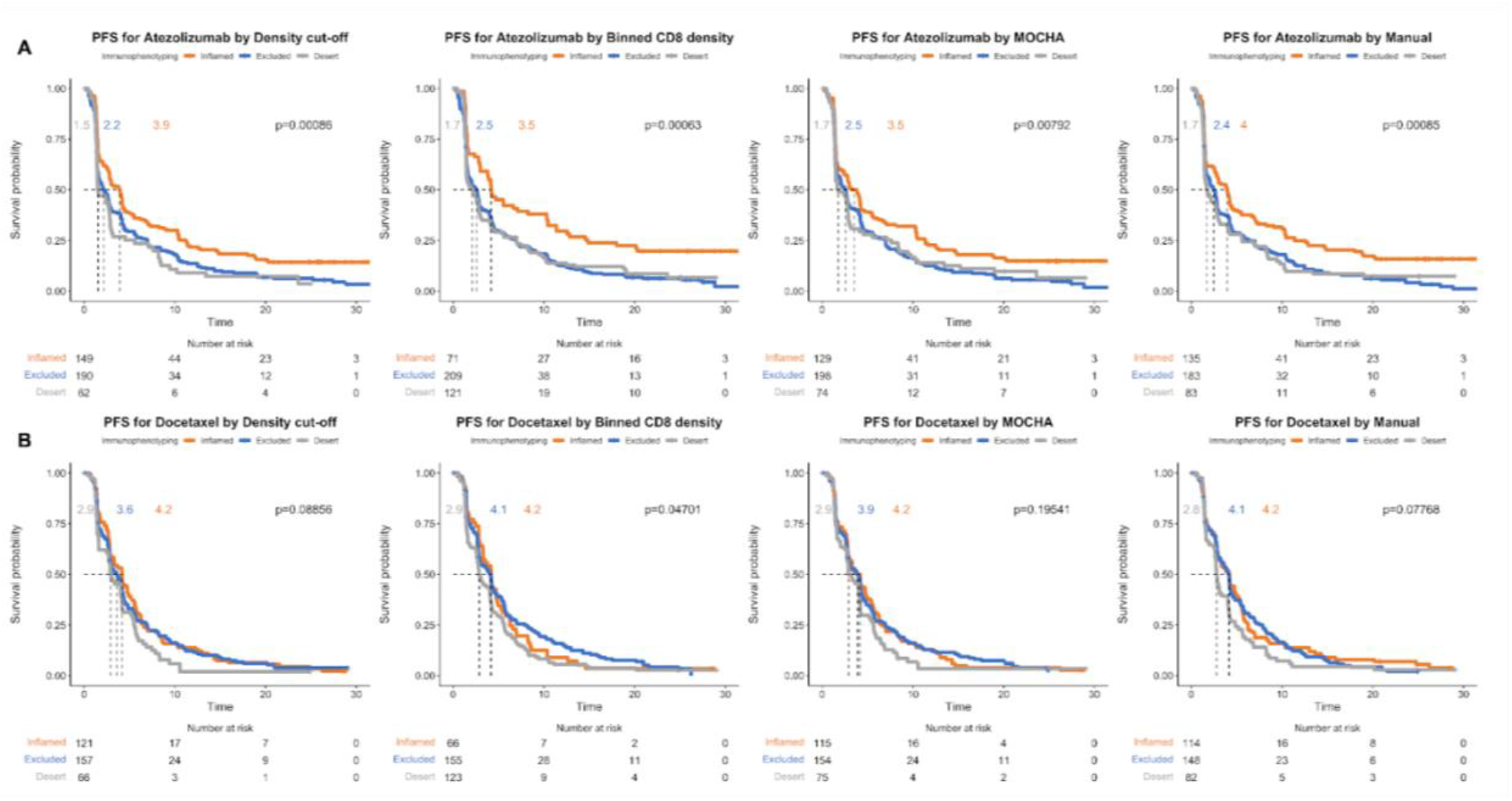
Kaplan–Meier curves for PFS in atezolizumab and docetaxel arms in OAK. Log-rank p-values are shown for tumor IP classes identified by the automated pipelines #1-3 and the manual method for atezolizumab arm (**A**) and docetaxel arm (**B**).

**Supplementary Figure 6.**
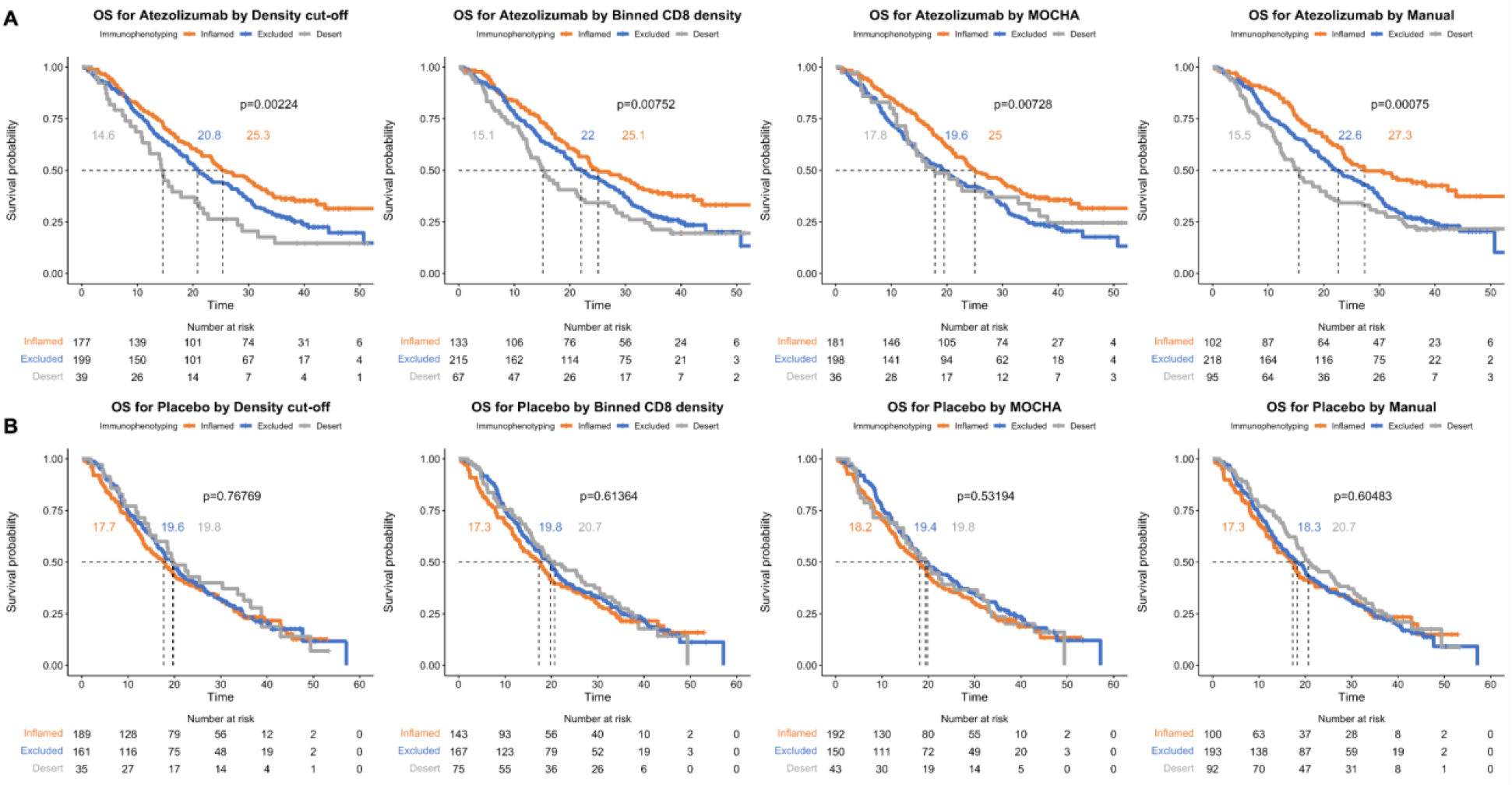
Kaplan–Meier curves for OS in atezolizumab and placebo arms in IMpassion130. Log-rank p-values are shown for tumor IP classes identified by the automated pipelines #1-3 and the manual method for atezolizumab arm (**A**) and placebo arm (**B**).

**Supplementary Figure 7.**
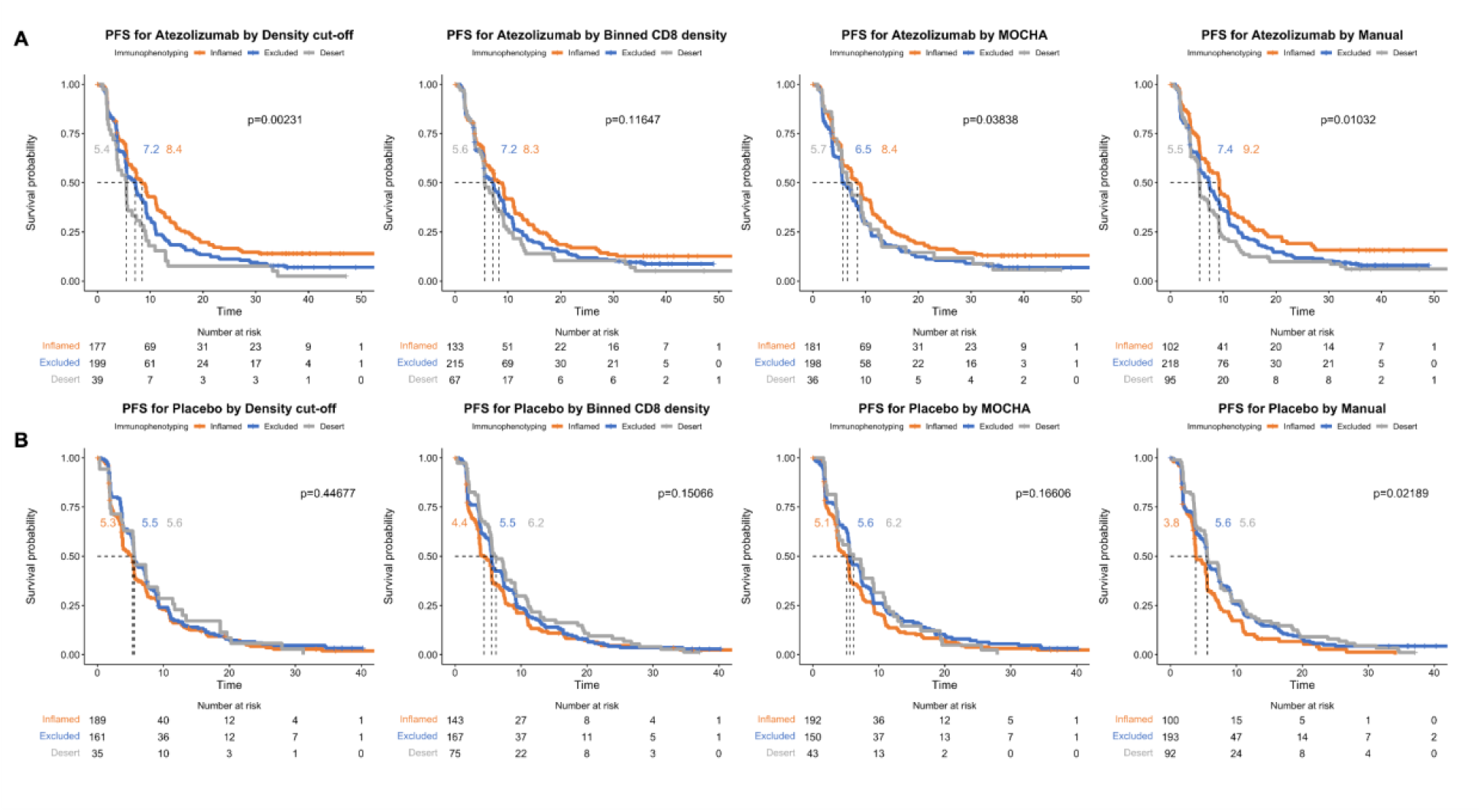
Kaplan–Meier curves for PFS in atezolizumab and placebo arms in IMpassion130. Log-rank p-values are shown for tumor IP classes identified by the automated pipelines #1-3 and the manual method for atezolizumab arm (**A**) and placebo arm (**B**).

